# Immunosuppression and senescence in dung beetles exposed to ivermectin

**DOI:** 10.1101/2025.09.14.676117

**Authors:** Sebastián Villada-Bedoya, Isaac G-Santoyo, Daniel González-Tokman

**Affiliations:** Neuroecology Lab, Facultad de Psicología, UNAM, Mexico City 04510, Mexico; Red de Ecoetología, Instituto de Ecología, A. C. Carretera Antigua a Coatepec 351, Xalapa 91073, Mexico

**Keywords:** aging, Coleoptera, ecotoxicology, immunoecology, macrocyclic lactones, Scarabaeinae

## Abstract

Immunosuppression and premature senescence are main risks of exposure to toxic compounds that might define individual longevity and the fate of natural animal populations. However, these sublethal effects have not been studied in insects that are of fundamental importance for human economy and well-being such as dung beetles. We exposed adult dung beetles *Euoniticellus intermedius* to ivermectin, a commonly used antiparasitic drug in cattle that is excreted in dung, and measured immune activity through phenoloxidase (PO) and its zymogen, prophenoloxidase (proPO), in males and females of different ages. We predicted that both ivermectin exposure and age would reduce immune activity, with the negative effect of ivermectin being more pronounced in older individuals, revealing immunosenescence. Despite PO and proPO activities decreased with age and ivermectin exposure, ivermectin effects on these immune mechanisms were constant across ages, revealing no immunosenescence caused by ivermectin. Whereas males suffered a reduction in PO and proPO with ivermectin and age, female proPO was not affected by ivermectin or age. This reveals a strategy of self-care adopted by females to prioritize immune function when facing stressful conditions. Among sublethal effects caused by ivermectin, immunosuppression might be a main physiological driver of population declines in contaminated environments.

## 1. Introduction

Senescent, old animals have a high probability of dying soon and will be less likely to mate as time passes [1], [2]. Despite senescence is unavoidable, for being the result of accumulated cellular damage with age, environmental factors experienced by individuals contribute to premature senescence [3]. Among the main causes of death in senescent animals is parasitism, as the immune system does not work as effectively as before, a process called immunosenescence [4]. The senescence of the immune system is also accelerated by past mating effort or exposure to stressful environmental conditions [5]. However, not all components of the insect immune system age similarly, as in *Tenebrio molitor* beetles, where enzymatic and cellular components but not antimicrobial peptides show signs of aging [6]. Similarly, males and females also differ in their immune responses and their senescence rates, with males being usually the sicker sex [7] and aging earlier (Bonduriansky et al. 2008). These differences result from differential interests in males and females, leading to females investing more resources in self-care (i.e., immune response) and males prioritizing reproduction over maintenance [7].

In insects, senescence occurs even in species with very short life cycle, such as *Drosophila melanogaster*, which has served as a model for aging [8]. Moreover, the mechanisms of insect senescence imply deterioration in DNA, just as in mammals, including humans [9]. In damselflies (Odonata), senescence seems to be mediated by hormones that distribute energetic resources towards different functions and may cause precocious death in insects, which might take different reproductive tactics to compensate [10]. However, whether exposure to contaminants accelerates senescence has not been explored, despite a clear relationship in the physiological mechanisms used to cope with poisons, pathogens and other stressors [11].

Dung beetles (Coleoptera: Scarabaeinae) are fundamental insects in forests and livestock pastures around the world, as they bury dung, contributing to soil fertilization, pest control and to reduce greenhouse gas emissions from dung [12], [13]. In cattle pastures, dung beetles are exposed to residues of veterinary medications in dung, which represents a threat because these contaminants may cause death of larvae developing in dung [14]. In particular, ivermectin is the most widely used antiparasitic drug in cattle in several regions of the world, for being highly effective against a wide range of parasites, but its residues are highly toxic for dung beetle larvae and are considered a main cause of population declines [14], [15]. Although adult beetles appear to be less sensitive to lethal doses of ivermectin, increasing evidence suggests that sublethal exposure may result in physiological stress, including oxidative imbalance and activation of heat shock proteins (HSP), which may divert metabolic resources away from immune pathways [16], [17]. Moreover, ivermectin has been shown to impair neuromuscular and olfactory functions in dung beetles, potentially disrupting pathogen detection and immune activation [18], [19]. Unexplored sublethal effects could include immunosuppression and senescence, which would imply higher deterioration with age [20].

In this study we evaluate if exposure to ivermectin during adulthood causes immunosuppression and immunosenescence in dung beetles *Euoniticellus intermedius*, a species that is abundant in ivermectin-contaminated pastures [21]–[23]. In particular, we measured the activity of the main enzyme from the insect innate immune system [24], phenoloxidase (PO), and its zymogen prophenoloxidase (proPO), in males and females of different age exposed to ivermectin. If ivermectin is immunosuppressive, we expect a reduction in PO and proPO activities in ivermectin-exposed individuals across different age categories. If ivermectin also triggers immunosenescence, we expect lower PO and proPO activities in old than in mature or young individuals.

## 2. Materials and methods

### 2.1. Study subject

This study was carried out with the dung beetle *Euoniticellus intermedius* (Reiche, 1848), an Ethiopian species that has been introduced in different continents and has become highly abundant in some cattle pastures around the world. This species is a tunneler, small-sized (8-13 mm) beetle, and is currently one of the most abundant dung beetles in eastern Mexico’s livestock systems [25], [26]. Whereas larval development takes three to four weeks in the laboratory, adult longevity goes up to 140 days but is drastically reduced by mating [27].

### 2.2. Insect rearing conditions

Beetles were collected in september 2021 at Medellín, Veracruz, Mexico (18° 58’19.37’’ N, 96° 04’51.43’’ W; 37 m asl) in a ranch where cattle are not treated with ivermectin. Thirty males and 30 females were collected and grown in randomly-formed pairs of a male and a female in 1 L plastic containers filled 80% with sterile sieved soil. Beetles were fed *ad libitum* with cow dung collected in Acajete, Veracruz, Mexico (19° 36’17.78’’ N, 97° 00’01.16’’ W), where ivermectin is not applied to cattle. The dung (humidity ≈ 87.84 ± 0.1%, pH = 7.0) was homogenized in an industrial mixer (Bathammex, Mexico) and frozen to eliminate parasites at -18 °C for at least 48 h before feeding the beetles. Breeding was carried out under controlled conditions (27.5 ± 0.4 °C, RH = 74 ± 2.5%, photoperiod 12L:12D).

### 2.3. Experimental design

Experimental beetles were acclimated to laboratory conditions for two generations during which they were fed *ad libitum* with cattle dung. In F2 generation, newly emerged beetles were placed in couples of a male and a female and were allocated to treatments that were exposed to ivermectin- or control-treated dung at different age categories: young (3-7 days old), mature (8-27 days old), and old (> 27 days old) (Fig. 1). These categories were chosen based on the reproductive biology of *E. intermedius*, which reaches sexual maturity at 4-5 days and has a median survival of around 30 days (González-Tokman, 2022).

**Fig. 1.**
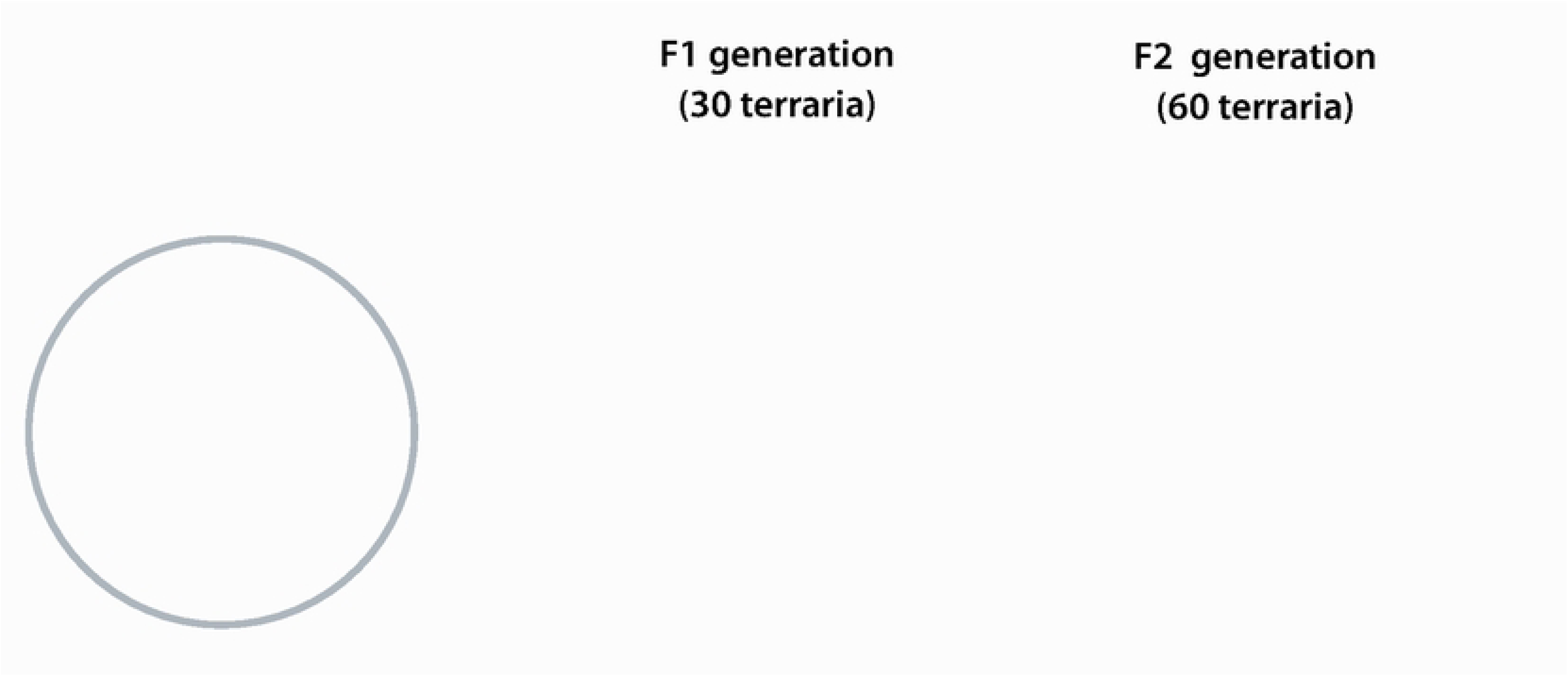
Experimental design, treatments and age categories.

To evaluate the effects of ivermectin across age categories, terraria for each age category were used to replace clean dung with ivermectin-treated dung (ivermectin treatment, see below). In these terraria, the dung was substituted five days before the established age was reached (Fig. 1). Beetles from the young age category and the ivermectin treatment were fed ivermectin-treated dung from day one. Once the required age was reached, the beetles were fasted for 24 h before measuring the variables of interest (PO and proPO, see below). Up to nine individuals from each terrarium were used to represent different age categories.

For the ivermectin treatment, we mixed the dung with ivermectin (CAS-Number: 70288-86-7, Sigma-Aldrich; purity of 90% ivermectin B_1a_ and 5% ivermectin B_1b_; lot number: SLBG8734V) diluted in acetone (CAS-Number: 1567-89-1, Sigma-Aldrich; purity > 99.8). We prepared the experimental treatment using a concentration of 10 µg.kg^-1^ (10 µg of ivermectin in 10 mL of acetone per kg of fresh dung). In our studied species, the used ivermectin concentration resembles the one excreted in dung by cattle that was treated after ca. 28 days and has known negative effects on ovary physiology and larval survival, leading to 20-50% reductions in offspring emergence [17], [28], [29]. Higher concentrations (e.g., 30 mg.kg^-1^) have more severe effects on beetle emergence whereas lower concentrations (e.g., 3 mg.kg^-1^) have slightly smaller effects [29]. Therefore, our used concentration represents a field-realistic, intermediate concentration which is suitable to test for sublethal effects and immune responses to ivermectin [16], [28]. The control treatment had 10 mL of acetone per kg of fresh dung. This vehicle-treated control has no negative effects on *E*. *intermedius* dung beetles compared to the intact control dung [17], [29]. Thawed dung already prepared with the treatment (i.e., with ivermectin or acetone) was always kept at 4 °C.

### 2.4. Measurement of phenoloxidase and prophenoloxidase

Most invertebrates exhibit melanization as a major innate immune response, initiated by the enzymatic conversion of the zymogen prophenoloxidase (proPO) into the functional enzyme phenoloxidase (PO). Phenoloxidase then catalyzes the conversion of phenols into quinones, which subsequently polymerize into melanin [30]. Due to this elemental function, PO and its zymogen proPO are important components of the insect immune response that have been linked to resistance against a wide variety of parasites and pathogens [31], [32]. PO is usually found as inactive zymogens, proPO, in all insects and is converted to active-PO when required by infection or wounding of the cuticle [33].

Synthesis of proPO occurs mostly in hemocytes [34] and is activated in response to foreign agents such as lipopolysaccharides, bacteria, fungi, damaged cells and toxic compounds [24]. As a result, when evaluating insect immunity, two distinct measurements of PO can be obtained: the enzyme that is readily available and naturally activated within the insect (PO activity) and the total investment in the PO pathway encompassing both proPO and active-PO (proPO activity), which is assessed after artificially activating the precursor proPO. To determine PO activity, one can directly measure it as described below. However, to measure proPO activity, proPO needs to be transformed into PO with the help of α-chymotrypsin (see below) [24].

A total of 247 beetles were used for measurement (PO = 218 inds; proPO = 213 inds). As far as possible, PO and proPO activity was quantified for each individual but in the end the number of individuals used for each measurement was limited by the amount of hemolymph that was extracted from each one. To measure PO and proPO, we extracted haemolymph from individuals and diluted in phosphate buffered saline (PBS, 1x; for each 1 µl of hemolymph, 15 µl of cold PBS were added). Subsequently, the samples were frozen at -18°C until the samples were analyzed under a microplate reader (BioTek ELX800). Each sample was used to quantify PO and proPO. To standardize enzyme activity by total protein in the haemolymph, protein content was also measured. To measure proteins, the bicinchoninic acid assay (BCA) was used. Briefly, 5 μl of haemolymph extract was added to a microplate well containing 45 μl of distilled water (ddH_2_O). Next, 150 μl of BCA solution (BCA Protein Assay Kit 23225; Thermo Scientific, Rockford, Illinois) was added. After 9 min of reaction at room temperature, the absorbance was measured at 590 nm, and the amount of protein relative to the albumin standard was obtained from the same kit.

The remaining extract was used to measure PO and proPO activities (see similar procedures in Busso et al., 2017; González-Santoyo et al., 2010). To measure PO activity, we added 10 μl of beetle extract to a microplate well containing 190 μl of L-Dopa solution (the substrate for the PO, Sigma D9628, 2 mg.ml^−1^ of ddH_2_O). Phenoloxidase activity was quantified at 28 °C, at an absorbance of 490 nm for 1 h. To measure proPO activity, 10 μl of beetle extract was added to a microplate well containing 3 μl of α-chemotrypsin (Sigma C7762, 2 mg.ml^−1^ of ddH_2_O) and left 10 min at room temperature to convert proPO into PO. Next, 27 μl of cold PBS and 140 μl of L-Dopa solution were added to the mix and the colorimetric reaction was followed in a microplate reader. Readings were taken for 4 h at 490 nm. Controls in both essays followed the same procedures but using cold PBS instead of beetle extract. To control for L-Dopa intrinsic conversion to dopaquinone, we subtracted control assay activity. Afterward, we standardized these values by dividing them by the protein amount in the sample. The reported values indicate activity per μg of protein (see similar procedure in Busso et al. 2017). In this study, each of our categories consisted of measurement of PO and proPO activity in adult dung beetles: **i)** measurement of PO in young (♀=41, ♂=38), mature (♀=36, ♂=27) and old beetles (♀=51, ♂=25), and **ii)** measurement of proPO in young (♀=35, ♂=37), mature (♀=39, ♂=31) and old beetles (♀=45, ♂=26).

### 2.6. Statistical analyses

All statistical analyses were performed in R program version 4.3.0 [37] using RStudio [38]. Type II analysis of variance tables based on 1- df chi-squared tests were used to assess significance in all models using the car library [39]. We summarized and organized the data using the dplyr library [40]. Figures were made using the ggplot2 library [41].

We compared PO and proPO activities for control and treated beetles across age categories using generalized linear mixed models (GLMMs) with the ‘glmer’ function of the lme4 package [42] and ‘gamma’ distribution, following Zuur et al. (2009). In these models, the response variables were PO or proPO activity, and the explanatory variables were: ivermectin treatment, sex, age, and the second- and third-degree interactions. The random effect in all GLMMs was the family (terrarium, considering that some samples came from siblings).

In all the above models, starting from the most parametrized model, we chose the most parsimonious model using the Akaike Information Criterion (AIC) [44]. We considered the most parsimonious model to be the one with the lowest AIC value and a ΔAIC greater than two units from the second most supported model [45]. When the difference between the most parsimonious model and the next model was Δ<2, we adhered to the statistical principle of parsimony, selecting the minimum adequate model with the lowest AIC and the fewest parameters [46]. Variables excluded by this criterion were considered non-significant for the model. For all analyses, model validation was done by inspecting normality of the residuals using Q-Q charts and homogeneity of variances through the Fligner-Killeen test [47]. Also, we assessed the presence of outliers using Cook’s distances.

## 3. Results

### 3.1. PO and proPO activities

In *E*. *intermedius*, PO and proPO activities were lower in ivermectin-exposed beetles compared to control-treated individuals (Table 1). As there was a treatment by sex interaction explaining both PO and proPO (Table 1), males and females were analyzed separately. In females, PO activity was reduced by ivermectin treatment and age, with mature and old females showing lower PO activity than young females (Fig. 2a). On the other hand, ivermectin had no effect on female proPO activation but young individuals showed higher proPO activation than mature and old females (Table 1; Fig. 2b).

**Fig. 2.**
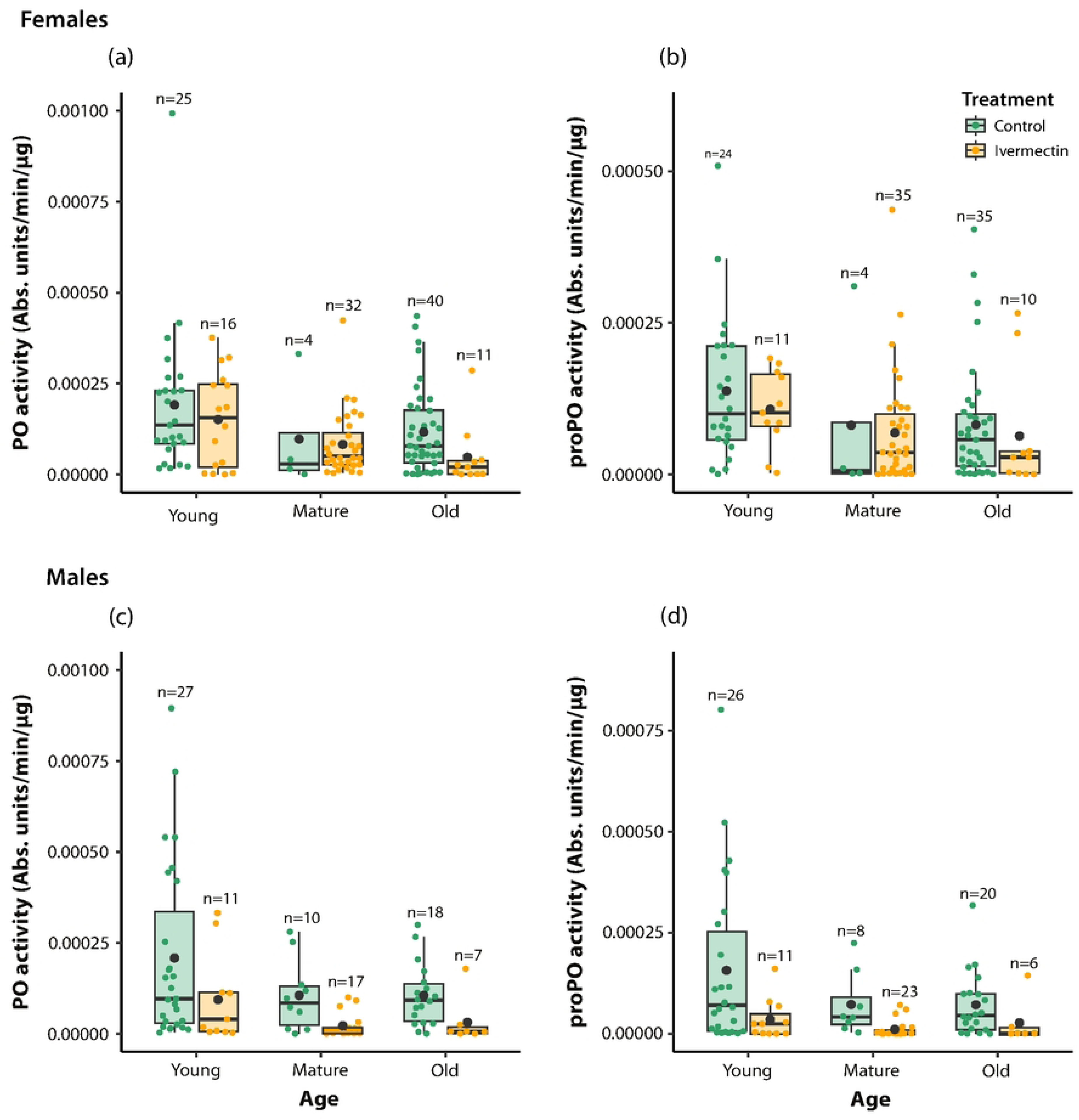
Effect of ivermectin treatment on phenoloxidase (PO) and prophenoloxidase (proPO) activities in females (a, b) and males (c, d) of *Euoniticellus intermedius* dung beetles of different age categories. Enzyme activity was measured as absorbance units/min/μg of protein. The line within each box represents the median, the boxes represent the first and third quartiles, and the whiskers represent the maximum and minimum values. The black dot represents the mean. Each colored dot represents a beetle.

**Table 1.** Results of generalized linear mixed models (GLMMs) analyzing the effects of ivermectin on male and female *Euoniticellus intermedius* beetles of different age categories on PO and proPO activities. The random effect was the family. Significant (* α < 0.05) and marginally significant (**·** α < 0.1) effects are shown in bold. Abbreviations: df, degrees of freedom; *χ*^2^, chi-squared; NS, effect not retained in the best supported model.

| Explanatory variables | df | Response variables |  |  |  |
| --- | --- | --- | --- | --- | --- |
|  |  | PO |  | proPO |  |
| | | $\chi^2$ | P | $\chi^2$ | P |
| Treatment | 1 | **15.49** | **<0.001\*** | **13.91** | **<0.001\*** |
| Sex | 1 | **5.06** | **0.024\*** | **11.55** | **0.001\*** |
| Age | 2 | **18.27** | **<0.001\*** | **10.88** | **0.004\*** |
| Treatment × Sex | 1 | **5.43** | **0.020\*** | **14.54** | **<0.001\*** |
| Treatment × Age | 2 | NS | NS | NS | NS |
| Sex × Age | 2 | NS | NS | NS | NS |
| Treatment × Sex × Age | 2 | NS | NS | NS | NS |
| **Females** |  |  |  |  |  |
| Treatment | 1 | **3.81** | **0.051•** | 0.64 | 0.422 |
| Age | 2 | **7.92** | **0.019\*** | 3.68 | 0.159 |
| Treatment × Age | 2 | 1.81 | 0.404 | 0.01 | 0.993 |
| **Males** |  |  |  |  |  |
| Treatment | 1 | **15.85** | **<0.001\*** | **18.89** | **<0.001\*** |
| Age | 2 | **11.88** | **0.003\*** | **8.29** | **0.016\*** |
| Treatment × Age | 2 | 1.23 | 0.541 | 1.03 | 0.598 |

In males, PO and proPO activities were lower in ivermectin-exposed than in control beetles (Table 1; Fig. 2c, d). Regarding the effects of age, young individuals had the highest values of PO and proPO, followed by old and mature males, which showed the lowest values (Fig. 2c, d). Nevertheless, the interaction treatment × age had no significant effect on male PO and proPO activities, revealing similar effects of ivermectin across ages in males (Table 1).

## 4. Discussion

In the present study, we found that ivermectin had immunosuppressive effects in adult dung beetles, adding evidence that this widely used antiparasitic drug [14] not only affects larvae, but also adults of these non-target insects of fundamental economic and ecological importance. Despite the measured immune responses declined with age, ivermectin effects on PO and proPO were consistent across age categories, in contrast to the predicted role of ivermectin in accelerating immunosenescence. The observed decline in immune responses with age might result from immunosenescence, but it could also result from physiological trade-offs involving allocation to other life-history traits, thereby affecting immunity [48]. Additionally, we acknowledge that other factors could contribute to senescence-like patterns like enviromental stressors (e.g., temperature fluctuations, habitat degradation), nutritional variation in dung, or genetic differences among individuals can modulate both immune function and aging trajectories [49]–[51]. Although we minimized such variables through standardized rearing conditions, future studies should investigate whether ivermectin acts synergistically with enviromental or dietary stress to accelerate functional decline, and whether genetic background modulates these effects. Experimental evidence challenging the insect immune system is still needed in dung beetles of different ages to determine whether immunosenescence or differential resource allocation are driving the observed effect.

Reductions in immune responses might be challenging for insects inhabiting pathogen-rich media such as dung [52], so that the consequences of ivermectin-caused immunosuppression might lead to rapid death in nature, with implications for population sizes and genetics [53]. Our results showing immunosuppressive effects of ivermectin in dung beetles are contrasting with evidence in dung flies *Scathophaga stercoraria* (Diptera: Scathophagidae) where PO activity was enhanced by ivermectin exposure [54]. However, the ivermectin concentration used in such study was 100 times lower than the dose used in the present study, indicating that the effects of ivermectin on insect immunity depend on the dose. Further studies evaluating a gradient of ivermectin doses should evaluate immune responses in coprophagous insects, also considering other immune effectors and pathways [55].

The immunosuppressive effects of ivermectin on dung beetles might arise through the negative effects that ivermectin has on different components of the host physiological condition and an integrated defense system which allows insects to cope with parasites and toxic compounds [11]. For example, evidence indicates that the ivermectin concentration used in the present study induced defensive physiological mechanisms such as production of heat shock proteins and antioxidants in our study species, also decreasing muscular mass in exposed individuals [16], [29]. As transcriptomic and proteomic analyses of ivermectin-exposed invertebrates indicate that mechanisms used in response to ivermectin include the detoxification metabolism, the mitochondrial respiratory chain and immune-related genes [20]; Gao et al. 2021; Álvarez-Sánchez et al. 2023), further studies in wild dung beetles from ivermectin-contaminated and ivermectin-free pastures could evaluate the immune responses and other physiological mechanisms to cope with ivermectin under challenging natural conditions, where pathogens and poisons simultaneously threat insect survival [11].

Despite reducing PO activity in females and males, ivermectin did not affect proPO in females as it did it in males. Sex differences in immune responses are common and result from different life-history strategies between the sexes, with females favoring maintenance, including immune responses, and males prioritizing reproduction, even if the cost is high in terms of survival [7], [56]. Despite males being more susceptible to infections than females, it remains to be tested whether it is a generality that males are also more sensitive to toxic compounds such as ivermectin. It also remains to be investigated why ivermectin-exposed females did not suffer changes in proPO as in PO, although the reasons must be associated to the rate of proPO conversion into PO or the multiple functions of proPO in insects beyond immunity, such as the formation of pigments based on melanin formation [57].

## 5. Conclusions

Immunosuppression and accelerated senescence are main effects caused by environmental stressors, with important consequences on individual survival and reproduction. Evidence presented here reveals that immune responses declined with age and ivermectin exposure in dung beetles, but ivermectin effects were constant across ages, suggesting no role of ivermectin in immunosenescence. Whereas ivermectin and aging led to immunosuppression in males, the effect on females was not as obvious. In a changing world where organisms face increasingly stressful conditions, individual immunosuppression and senescence can have important consequences at the population level for insects of fundamental importance for human well-being, with males and females showing different responses.

## Acknowledgements

Thanks to Dr. Imelda Martínez for logistic support. To R. Madrigal-Chavero for fieldwork help. We also thank Y. Gil-Pérez for her help in the laboratory. To the Consejo Nacional de Humanidades, Ciencias y Tecnologías (CONAHCYT) for the scholarship granted to S.V.-B. (CUV: 865045, Scholarship holder number: 491448) to carry out doctoral studies. The present study was funded by Consejo Nacional de Ciencia y Tecnología (CONACYT project Ciencia Básica 257894) granted to D.G.-T. The authors declare that no conflict of interest exists. We also thank to UNAM Posdoctoral Program (POSDOC), because during S.V.-B.’s postdoctoral stay it was possible to carry out this work in parallel to his stay in the Neuroecology Lab.

## Author contributions

**Conceptualization:** Daniel González-Tokman, Sebastián Villada-Bedoya.

**Data curation:** Sebastián Villada-Bedoya.

**Formal analysis:** Sebastián Villada-Bedoya, Daniel González-Tokman.

**Funding acquisition:** Daniel González-Tokman.

**Investigation:** Daniel González-Tokman, Sebastián Villada-Bedoya, Isaac G.-Santoyo.

**Methodology:** Daniel González-Tokman, Sebastián Villada-Bedoya.

**Project administration:** Daniel González-Tokman.

**Resources:** Daniel González-Tokman.

**Writing – original draft:** Sebastián Villada-Bedoya, Daniel González-Tokman, Isaac G.-Santoyo.

**Writing – review & editing:** Sebastián Villada-Bedoya, Daniel González-Tokman, Isaac G.-Santoyo.

**Competing interests:** The authors have declared that no competing interests exist.

